# Predicting the potency of anti-Alzheimer drug combinations using machine learning

**DOI:** 10.1101/2020.04.28.066340

**Authors:** Thomas J Anastasio

## Abstract

**BACKGROUND:** Clinical trials of single drugs for the treatment of Alzheimer Disease (AD) have been notoriously unsuccessful. Combinations of repurposed drugs could provide effective treatments for AD. The challenge is to identify potentially potent combinations.

**OBJECTIVE:** To use machine learning (ML) to extract the knowledge from two leading AD databases, and then use the machine to predict which combinations of the drugs in common between the two databases would be the most effective as treatments for AD.

**METHODS:** Three-layered neural networks (NNs) having compound, gated units in their internal layer were trained using ML to predict the cognitive scores of participants in either database, given the other data fields including age, demographic variables, comorbidities, and drugs taken.

**RESULTS:** The predictions from the separately trained NNs were strongly correlated. The best drug combinations, jointed determined from both sets of predictions, were high in NSAID, anticoagulant, lipid-lowering, and antihypertensive drugs, and female hormones.

**CONCLUSION:** The results suggest that AD, as a multifactorial disorder, could be effectively treated using a combination of repurposed drugs.

## BACKGROUND

Artificial intelligence (AI) and machine learning (ML) hold great promise for advancing biomedicine [1-3]. Meanwhile, the development of effective treatments for Alzheimer disease (AD) and other neurodegenerative disorders is lagging that of other major diseases 4-6]. Consequently, increasing numbers of people in developed countries are living long enough to become demented. The power of AI and ML must be brought to bear on the pressing problem of AD.

Interest is growing in the AD community for using combinations of repurposed drugs as AD treatments 7-11]. The main challenge is to identify the best candidates from among the many possible combinations. AI and ML are uniquely well suited to address this challenge. They could be used to leverage the knowledge contained in AD databases to make the best attainable guess as to which combinations of repurposed drugs could be used to treat AD.

The goal of this work is to train AIs using ML to extract the knowledge contained in two, leading AD databases, and use them to identify drug combinations that could be effective in treating AD. The two AD datasets were obtained from the Rush Alzheimer Disease Center (RADC) and the National Alzheimer Coordinating Center (NACC). Both databases are highly regarded and used worldwide. NACC encompasses the data from the 29 Alzheimer Disease Centers (ADCs) that are funded by the National Institute on Aging of the National Institutes of Health of the United States [12,13]. RADC is an ADC but most of its dataset comes from the participants in the Religious Orders Study and Rush Memory and Aging Project (ROSMAP) [14,15]. A small amount of legacy RADC data is included in NACC (less than 1.4% of total NACC data). All RADC data was removed from the NACC dataset for this study. The two datasets used for ML in this study are entirely independent of one another (see Supplementary Note N1).

Neural networks (NNs) are the state-of-the-art AIs for generalizing from available data and making predictions. NNs can be configured in many different ways. Evaluation of a large number of possible configurations showed that the NNs that were optimal for the ROSMAP or NACC datasets were essentially the same. As a crosscheck, NNs having the same configuration were trained separately on the ROSMAP or NACC datasets and their predictions concerning the potency of many drug combinations were compared. The strong correlation between the two sets of predictions increases confidence in the jointly determined, best drug combinations.

## METHODS

Neural networks (NNs) are computational devices that accept an input on their input units, process it using internal units, and produce an output on their output units (see Figure 1). The ML technique used to train NNs is supervised learning, which requires a dataset composed of many input/desired-output examples. In supervised learning, an input is processed to form an output, and the actual output is compared to the desired output to form an error. The error is used to modify the weights of the connections between the units in the network, in such a way that the error between the actual and desired output for that input is reduced.

**Figure 1.**
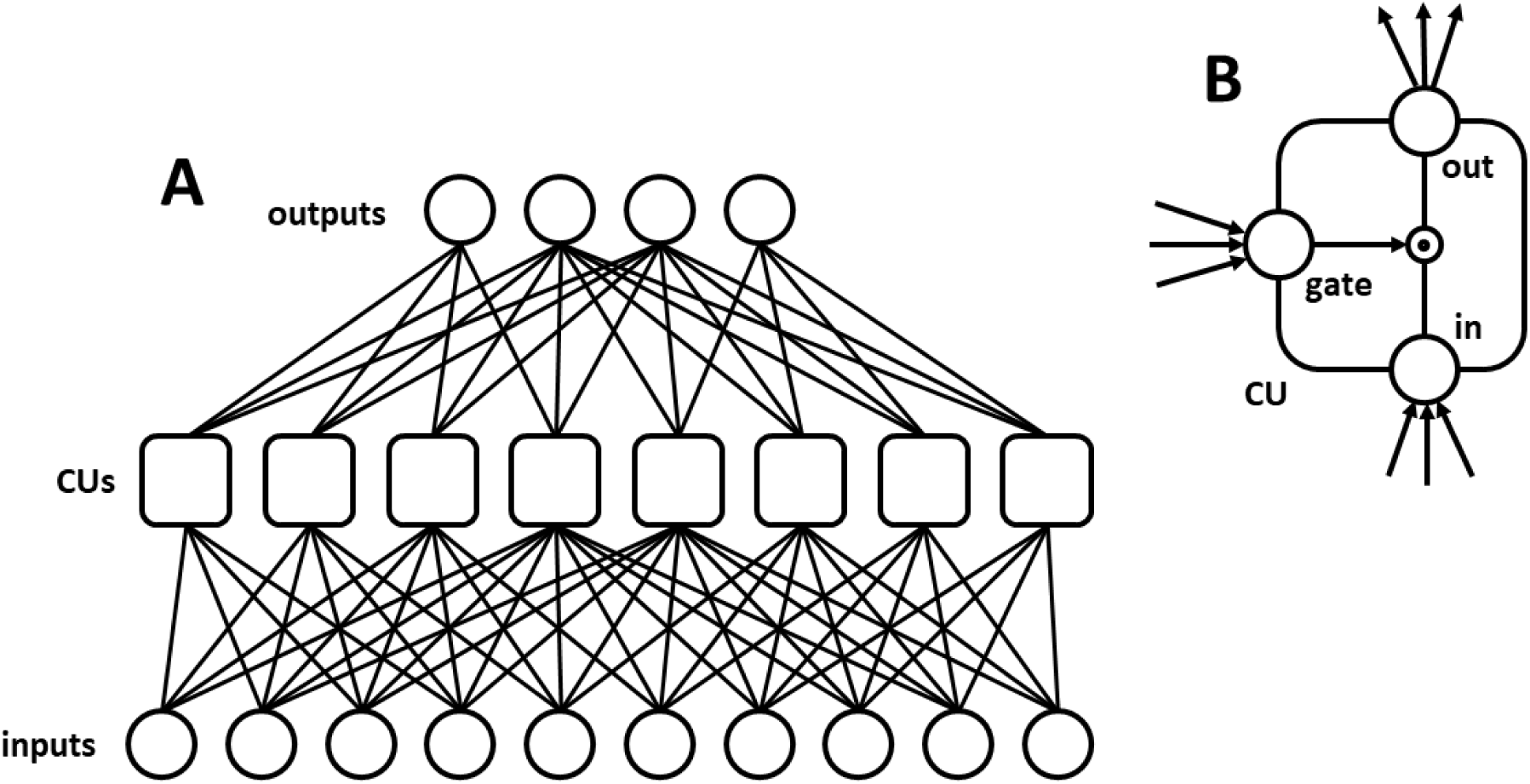
Drug combination potencies were predicted using trained NNs composed of simple and compound units. (A) The NNs had three layers: input, internal, and output. All input units connected to all internal units, and all internal units connected to all output units. NNs trained on the ROSMAP dataset had 113 input and 25 output units, while NNs trained on the NACC dataset had 101 input and 57 output units. Both NNs had 80 internal units. (B) The internal units were compound units (CUs). Each CU had its own input, output, and gate unit. Both the input and the gate unit of each CU received connections from input-layer units. The gate unit modulated the activation of the CU input unit before it was sent to the CU output unit. Each CU output unit connected to every output-layer unit.

The NNs considered here had three layers: input, output, and internal. Forward connections connected input to internal units and internal to output units. The input layer was composed of linear units that each represented the value of an input variable, scaled into the range [0,1]. The output layer was composed of nonlinear units that computed the sum of their weighted connections from internal units and imposed a sigmoidal activation function on it, squashing it into the range [0,1]. The numbers of units in the input and output layers are fixed by the dataset on which the NN is trained. The number of possible configurations of a three-layered NN can be increased at its internal layer.

Compound units known as long short-term memory units (LSTMs) composed the internal layer. LSTMs encompass several nonlinear units that interact within each LSTM [16,17] (see Supplementary Note N2). There is no limit on the number of internal LSTMs in an NN, and they can receive forward connections from input units as well as recurrent connections from each other. The best configuration for a given application may be a simplification of a recurrent network of LSTMs.

Recurrent LSTM networks are trained using an ML algorithm known as backpropagation through time (BPTT) [18,19]. BPTT draws temporal sequences of input/desired-output examples from the dataset at random (with replacement), and continues training the NN over many iterations, thereby reducing the error over the dataset. The version of BPTT used here has three parameters: starting and ending learning rate and fixed momentum. The learning rate decreases from starting to ending as the training iterations progress from first to last, and the momentum determines how much of the previous iteration’s learning should be applied on the current iteration. These learning parameters will likely vary between NNs having different configurations.

Once trained, the NN not only can reproduce the desired outputs for the inputs in the dataset but can also generalize the knowledge it has extracted, and can predict the outputs for inputs on which it has not been trained. To assess generalizability, the dataset is divided into a training set (usually 75% of the dataset) and a testing set (the other 25%). Then the NN is trained on the training set and tested on the testing set. The generalization error is the total error between the actual (predicted) and desired outputs over the testing set.

BPTT entails two sources of randomness: random initial connection weights and random presentation of input/desired-output sequences. Consequently, NNs trained on the same dataset can vary in their outputs, and the best predictions are derived from the averaged outputs of many trained NNs [20]. Generalizability, which is essentially the ability of an NN to predict desired outputs over the testing set, is likewise best assessed by averaging the actual outputs of many retrained NNs.

The overall goal of this work was achieved in three steps. The first was to process the data contained in two AD databases into a form suitable for training NNs using ML. The second step was to choose the best NN configuration for this application. The third step was to use the best NN to perform the computational drug-combination screens that provided the results of this study.

### Processing the data from two AD databases

The RADC and NACC databases had similar fields which included, for each participant, age at visit, demographics (sex, race, etc), tobacco and alcohol use, comorbidities (cancer, stroke, etc), drugs taken, and the results over a battery of cognitive tests. The data in these fields were arranged into input/desired-output pairs, where the desired outputs were the cognitive test results and the inputs were the values for all the other fields for each visit. Because age is a key variable, all of the input/desired-output pairs were arranged into age-advancing sequences for each participant.

Clinical databases entries are often incomplete. For the development of an NN training set, missing data cannot simply be assigned a value of zero. Missing input data was replaced by the average of the data for that field. Missing desired-output data was left blank, but during training all errors associated with missing desired-output data were set to zero. Because connection weight modifications are based on error, no weight modifications were made due to missing desired-output data.

Redundant database fields were removed. Ideally, the values assigned to an input or desired output should increase monotonically with the variable it represents, but database field codes can deviate widely from that simple relationship. Specific inputs or desired-output values were rearranged or reassigned, when necessary, to bring them all into a monotonically increasing regime. Finally, the input and desired-output data (except for missing desired-output data) were normalized to [0,1].

### Choosing the best NN configuration

The combination of NN features and ML parameters providing the best generalizability was determined using the genetic algorithm (GA) [21,22] (see Supplementary Note N3). Because of the randomness inherent in the GA, the best solution represents the consensus over multiple GA runs. Ten GA runs were performed separately for the ROSMAP and NACC datasets, producing twenty runs altogether. The GA results were similar between the two datasets and the consensus was obvious (see Supplementary Table T1). Using GA optimization, the best NN for this application lacked recurrent connections, and had LSTMs simplified to a single gating interaction in its internal layer (see Figure 1).

The use of gating is common in NN applications in which different internal units learn to specialize for different subsets of the dataset [23]. The fact that the consensus NN employed gated units suggests that such specialization improved generalizability in this application. The consensus NN had about 80 gated units in its internal layer, and its optimal ML parameters were 0.0600, 0.0006, and 0.0002 for starting and ending learning rate and fixed momentum, respectively.

### Performing the computational drug-combination screens

Consensus NNs were trained on the ROSMAP or NACC datasets separately, because the RADC and NACC databases have many structural differences and they cannot be merged without making many unsupportable assumptions. The trained NNs were used to screen all possible combinations of the drugs in common between the two databases. The drugs in both databases were listed in drug categories rather than as specific drugs. The categories were: non-steroidal anti-inflammatory drugs (NSAIDs, mainly cyclooxygenase-2 (COX2) inhibitors and aspirin); angiotensin converting enzyme (ACE) inhibitors; angiotensin II receptor blockers (ARBs); calcium channel blockers; beta blockers; diuretics; vasodilators; anticoagulants; lipid-lowering drugs; antidiabetics; estrogen or progestin; anti-adrenergics; antidepressants; antipsychotics; anxiolytics; anti-Parkinson drugs; and anti-Alzheimer drugs (mainly acetylcholinesterase inhibitors and N-methyl-D-aspartate receptor blockers). There are 131,072 combinations of these 17 drug categories, and consensus NNs, trained on either the ROSMAP or NACC datasets, were tested on every combination. This constituted two, separate computational screens (ROSMAP or NACC) over all 100K+ drug combinations.

The inputs to the NNs were all data fields other than cognitive scores. They included demographics, tobacco and alcohol use, and comorbidities as well as drugs taken. Valid comparison of the predicted potencies of different drug combinations required testing each drug combination using a standard input, in which all inputs besides drugs were the same. The standard input was based on an age-advancing sequence of 100 ages from 50 to 110 years, which encompasses the age range of the participants in the two databases. All of the other inputs, separately for either database, were resampled to conform to this sequence by computing a value that is representative of each input field at each age in the standard sequence (see Supplementary Note N4).

Then a consensus NN had its weights randomized and was trained on a full dataset (either ROSMAP or NACC), and the cognitive scores (represented by NN output units) predicted by the trained NN for each drug combination at each age in the standard age-advancing sequence were found. This process (randomization, retraining, and predicting) was repeated 100 times for either dataset, and the predicted cognitive scores were averaged over the 100 NNs. A combined cognitive score for each age in the standard sequence was computed by averaging over the individual predicted cognitive scores (represented by individual NN output units). The combined cognitive scores as predicted by one NN for many representative (not all 100K+) drug combinations versus age are shown in Figure 2.

**Figure 2.**
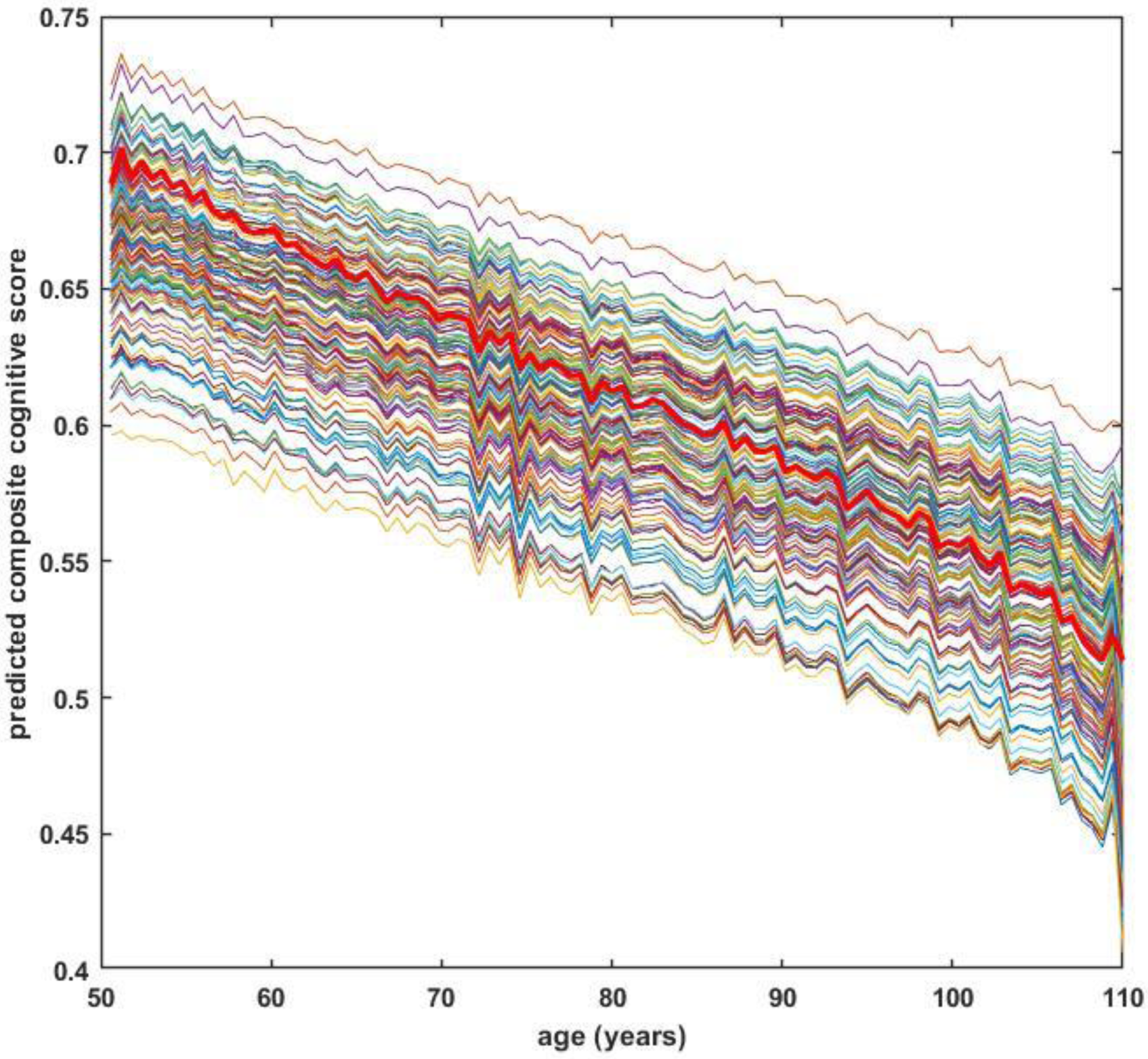
The combined cognitive scores as predicted by a single NN for each age in the age-advancing sequence for 65 representative drug combinations (every 2000^th^ combination selected from the full set of 131,072 combinations of 17 drugs). Each output unit represented the score of a different cognitive test, so the combined cognitive score was the average over the output unit activations. The combined cognitive score in the no-drug case is shown as a heavy red line. This NN was trained on the ROSMAP dataset. The results for NNs trained on the NACC dataset are similar (see Supplementary Figure S1).

Figure 2 shows that the combined cognitive score, which is the NN’s prediction of cognitive ability at each age for a specific drug combination, can be better than the no-drug case (heavy read line) for many drug combinations. To obtain a single number for the predicted potency of each combination, the differences between the combined cognitive scores predicted for that drug combination and the cognitive scores predicted for no drugs at all ages in the age-advancing sequence were averaged. The results of this study are reported in terms of the potency of each specific drug combination, as predicted from consensus NNs trained either on the ROSMAP or NACC datasets. For brevity, these separately trained NN predictions will be referred to as ROSMAP predictions or NACC predictions.

## RESULTS

The results presented in Figure 2 show that the combined cognitive scores predicted from NNs can be greater for many drug combinations than for no drugs. The scores shown in Figure 2 were predicted from NNs trained on the ROSMAP dataset; those predicted from NNs trained on the NACC dataset are similar (see Supplementary Figure S1). The ROSMAP or NACC predictions could be taken at face value, but our confidence in the two sets of predictions would be strengthened if they agreed. Figure 3A shows that they do agree (see also Supplementary Figure S2).

**Figure 3.**
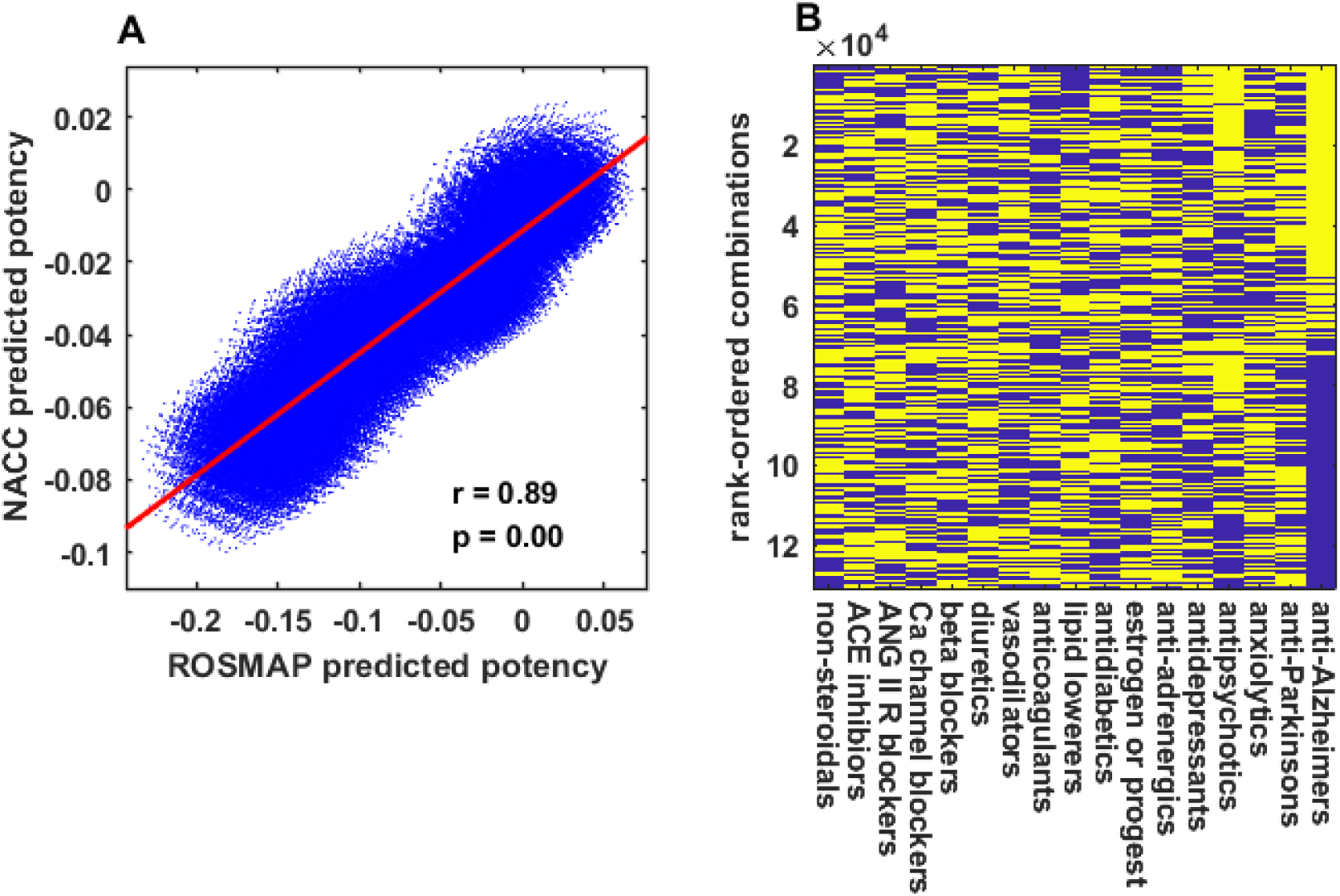
Predicting potent drug combinations jointly from NNs trained separately on the ROSMAP or NACC datasets for the 131,072 combinations of 17 drugs. (A) The potencies predicted by NNs trained on the ROSMAP or NACC datasets are strongly correlated. Each blue dot represents one drug combination, located by its ROSMAP and NACC predicted potency (r is the correlation coefficient, p is the probability that the correlation occurred by chance). (B) All 131,072 drug combinations are ranked according to predicted potency (top is highest) and displayed as a heat map (yellow, drug present; blue, drug absent). The most potent drug combinations include anti-Alzheimer drugs.

### Correlating ROSMAP and NACC drug-combination predicted potencies

The strong correlation between the ROSMAP and NACC predictions motivates joint predictions based on both dataset together rather than on either one alone. Drug combination potency can be predicted jointly from the ROSMAP and NACC predictions in terms of the value of the projections of the drug combination points onto the regression line. All 131,072 combinations of the 17 drug categories, rank ordered according to their jointly predicted potency, are shown as an image in Figure 3B.

From Figure 3B its seems that the benefit of a drug combination depends on whether or not it includes anti-Alzheimer drugs, whose primary effect is to improve the cognitive ability of AD sufferers. Other drugs that affect mood and/or cognitive ability, especially antipsychotic drugs, also seem determinative of drug combination benefit (see also Supplementary Figure S3). It is possible that drugs that act on the brain and/or psychological level and that improve cognition, either directly or by improving mood or behavioral control, may mask the neuroprotective effects of other drugs.

### Restricting the drug-combination screens

Of interest here are drug combinations that may treat the underlying cause of AD, which is neurodegeneration. To uncover such potentially potent AD treatments, the drug combination screen was repeated following the removal of all drugs that act mainly on the brain and/or psychological level. Those included the anti-Alzheimer and anti-Parkinson drugs as well as the anti-adrenergics, antidepressants, antipsychotics, and anxiolytics. This left drugs in eleven categories. The ROSMAP and NACC predictions for their 2048 combinations were also strongly correlated, as shown in Figure 4A.

**Figure 4.**
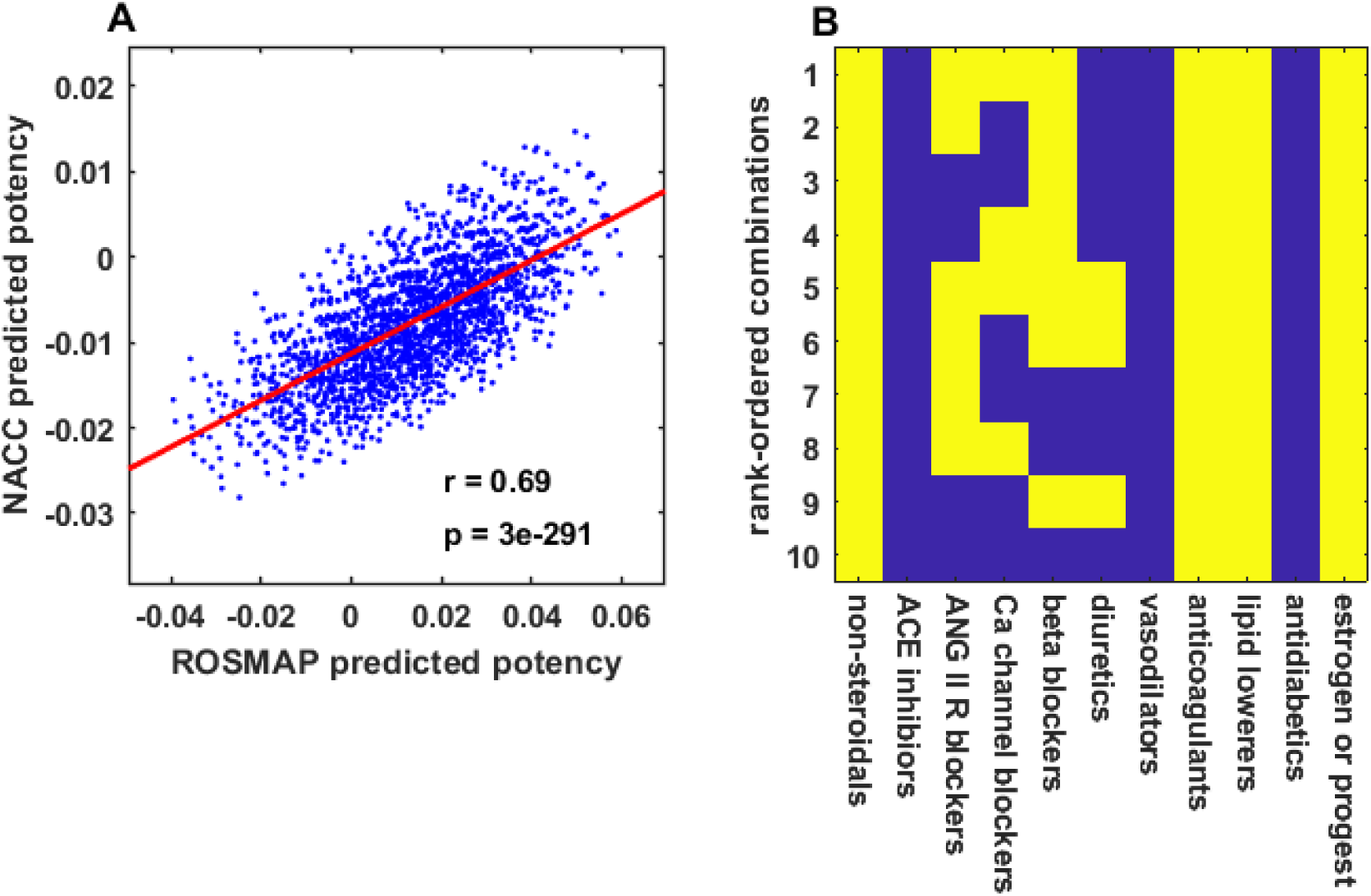
Predicting potent drug combinations jointly from NNs trained on the ROSMAP or NACC datasets for the 2048 combinations of the 11 drugs that do not target cognitive ability or mood. (A) The potencies predicted by NNs trained on the ROSMAP or NACC datasets are strongly correlated. (B) The top-ten drug combinations are ranked according to predicted potency (1 is highest). They include NSAID, anti-hypertensive, anticoagulant, and lipid-lowering drugs, and estrogen/progestin.

As shown in Figure 4B, the top-ten combinations of these eleven drug categories, jointly determined from the ROSMAP and NACC predictions, all included NSAID, anticoagulant, and lipid-lowering drugs, and the female hormones estrogen and/or progestin. The top nine also included one or more of these antihypertensive drugs: angiotensin II receptor blockers, calcium channel blockers, beta blockers, and diuretics.

While some men take estrogen or progestin medicinally, taking those hormones would not be appropriate for all AD sufferers. For that reason, the drug combination screen was repeated following the removal of estrogen/progestin, as well as all drugs that act mainly on the brain and/or psychological level. This left drugs in ten categories. The ROSMAP and NACC predictions for their 1024 combinations were also strongly correlated, as shown in Figure 5A.

**Figure 5.**
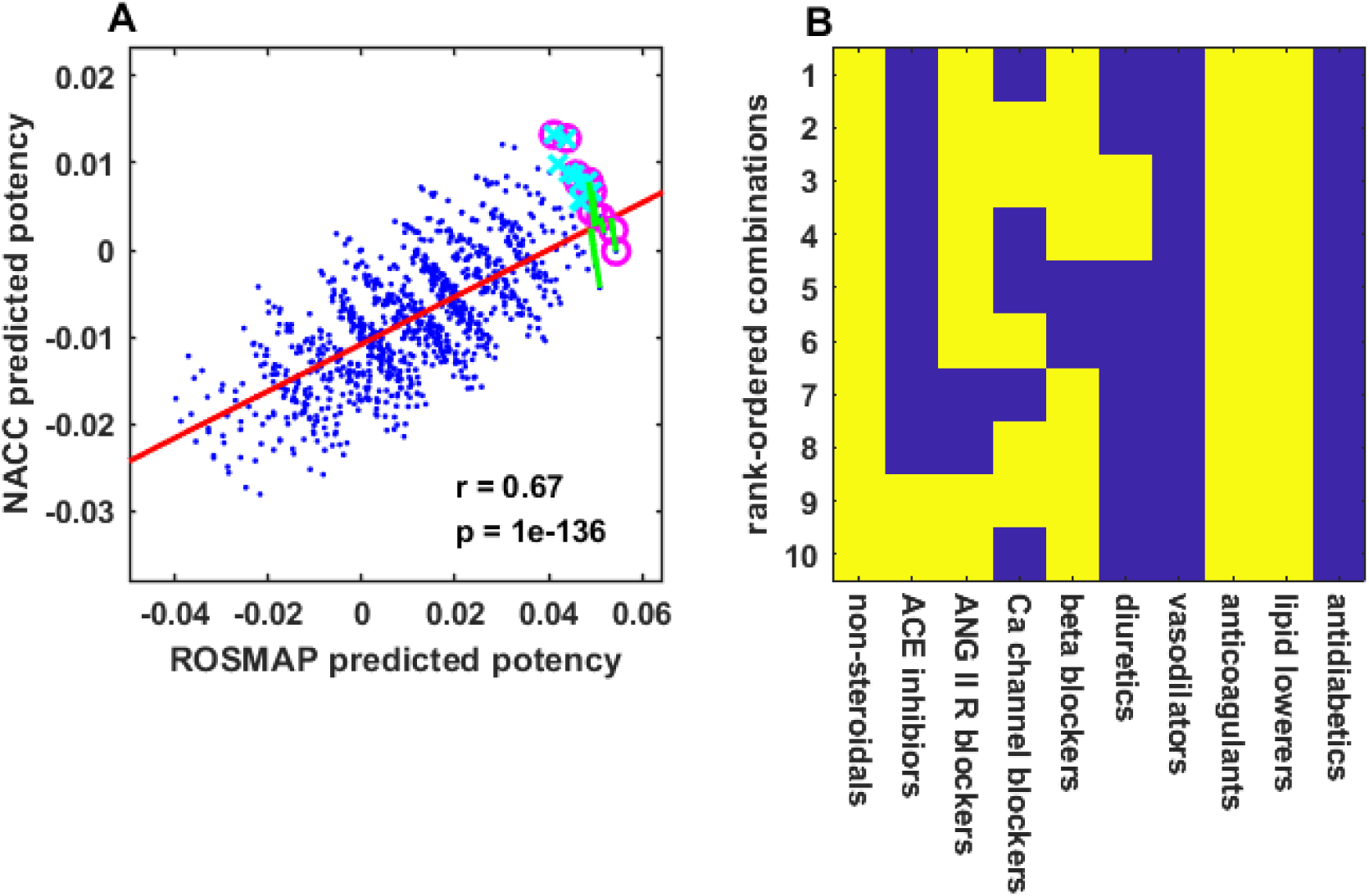
Predicting potent drug combinations jointly from NNs trained on the ROSMAP or NACC datasets for the 1024 combinations of the 10 drugs that do not include estrogen/progestin or that target cognitive ability or mood. (A) The potencies predicted by NNs trained on the ROSMAP or NACC datasets are strongly correlated. The ten-best drug combinations as determined jointly in terms of linear regression, sum of potency, or sum of rank of potency as predicted by NNs trained separately on the ROSMAP or NACC datasets are shown as green line segments, magenta o’s, or cyan x’s, respectively. (B) The top-ten drug combinations are ranked according to predicted potency (1 is highest). They include NSAID, antihypertensive, anticoagulant, and lipid-lowering drugs.

As shown in Figure 5B, the top ten combinations of these ten drug categories, jointly determined from the ROSMAP and NACC predictions, are similar to those that included estrogen/progestin. They all included NSAID, anticoagulant, and lipid-lowering drugs. Additionally, the top-ten combinations all included one or more drugs from all of the antihypertensive categories: ACE inhibitors, angiotensin II receptor blockers, calcium channel blockers, beta blockers, and diuretics.

### Alternative rankings of the best combinations

The simplest way to rank the drug combinations is separately in terms of the ROSMAP or NACC predictions alone. The top-ten drug combinations determined from ROSMAP or NACC predictions alone are similar to the jointly determined combinations in that they all include NSAID and lipid-lowering drugs, and most of them also include anticoagulant and antihypertensive drugs (see Supplementary Figure S4). Owing to the strong correlation between the ROSMAP and NACC predictions, the uncertainty associated with rankings based on ROSMAP or NACC predictions alone can be reduced by rankings that consider both sets of predictions together.

In previous sections, the joint determination was made in terms of the projections of the ROSMAP and NACC prediction points onto the line resulting from the regression of the NACC onto the ROSMAP predictions (NACC(ROSMAP)) (Figures 3A, 4A, and 5A). Because correlation is symmetric, rankings based on NACC(ROSMAP) or ROSMAP(NACC) regressions are identical (see Supplementary Figure S5). Joint rankings in terms of linear regression proceed most naturally from correlation analysis.

Other methods of jointly determining drug combination potency include ranking the sums of the ROSMAP and NACC predictions for each combination, or ranking the sums of the rankings of the combinations based on the ROSMAP or NACC predictions alone. The sets of ten-best drug combinations determined according to the sum or rank-sum methods overlap the set determined according linear regression, and encompass the points at the top-right of the scatter plot (see Figure 5A).

As shown in Figure 6, the top-ten combinations of the ten drug categories that exclude estrogen/progestin and the drugs that act mainly on the brain and/or psychological level, jointly determined using the sum, rank-sum, or linear regression methods are similar. They all included NSAID, anticoagulant, and lipid-lowering drugs. Additionally, most of the top-ten combinations included multiple antihypertensive drugs. The main difference was that the sum and rank-sum methods excluded ACE inhibitors while the linear regression method included them.

**Figure 6.**
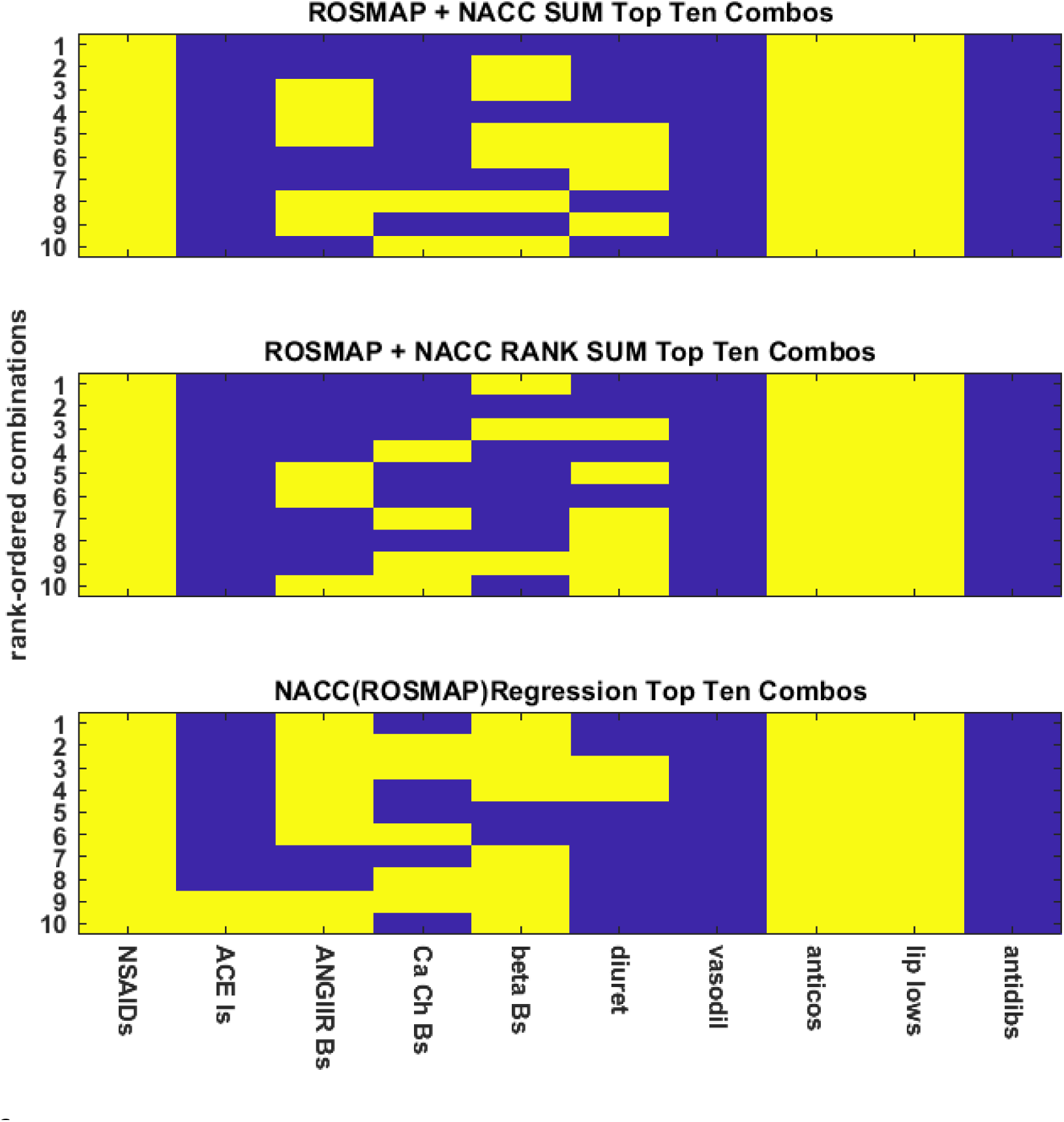
Different joint ranking methods yield similar orderings of drug combinations. The ten-best combinations of the ten drugs that do not include estrogen/progestin or that target cognitive ability or mood, as predicted jointly by NNs trained separately on the ROSMAP or NACC datasets all include NSAID, antihypertensive, anticoagulant, and lipid-lowering drugs whether the ranking is based on the sum of the separately predicted potencies (ROSMAP + NACC SUM), the sum of the ranks of the separately predicted potencies (ROSMAP + NACC RANK SUM), or the projections onto the regression line giving NACC potency as a function of ROSMAP potency (NACC(ROSMAP)). Note that the NACC(ROSMAP) and ROSMAP(NACC) regressions yield identical orderings of drug-combination predicted potency (see Supplementary Figure S5).

## DISCUSSION

However they are ranked, the ten-best combinations of the ten drugs shown in Figure 6, jointly determined from the ROSMAP and NACC predictions, include from three to seven drugs. This finding supports the general idea that multiple drugs may indeed be effectively used to treat neurodegeneration, which is a multifactorial disease[7-11]. The ten-best combinations are not random but tend to include drugs from the same categories: NSAID, anticoagulant, lipid-lowering, and antihypertensive drugs.

One possible interpretation of the results is that elderly individuals who take these drugs have better cognitive function than those who do not, because the conditions these drugs treat are associated with better cognitive function. This interpretation is ruled out by statistical analysis of the ROSMAP and NACC datasets showing that comorbidities, singly or in combination, do not improve cognitive function (see Supplementary Figures S6 and S7). The results are agnostic concerning mechanisms of action, which could involve both the primary and secondary effects of the drugs in the ten-best combinations. For example, certain NSAIDs (mainly COX2 inhibitors) not only reduce inflammation but also reduce amyloid-β production [24-26], while certain antihypertensive drugs also have anti-inflammatory properties [27-29].

The drugs that appear in the ten-best combinations in this study overlap those in a previous computational study in which ML was used to train a computational model of brain inflammation [30]. Model predictions of drug-combination efficacy were significantly correlated with the cognitive benefits of the same drug combinations as determined from the ROSMAP dataset. The ten-best combinations from the previous study also included NSAIDs (COX2 inhibitors and aspirin) as well as antihypertensive drugs (ACE inhibitors, angiotensin II receptor blockers, calcium channel blockers, and beta blockers). Earlier computational studies of potential anti-AD drug interactions were based on process-driven models, rather than on data-driven models as trained by ML. Those models suggested that NSAIDs could be effectively combined with antidiabetic drugs or estrogen as anti-AD therapies [31,31]. The drug categories in the previous studies were not the same as in this study. That discrepancy limits the value of directly comparing their results.

The ten-best combinations of the eleven drugs that excluded drugs acting on the brain and/or psychological level included NSAID, anticoagulant, lipid-lowering, and antihypertensive drugs, as well as estrogen/progestin. Drugs from all of these categories have been shown to reduce AD risk, however they all show disappointing results when used singly in clinical trials [33-45]. The use of combinations of these and other drugs remains an attractive option [7-11].

All drugs in the computational combination screens in this study are approved. Many elderly already take some of these drugs. Where not contraindicated, other drugs could be added to their regimens to complete some of the predicted combinations, for the purpose of conducting controlled clinical trials to test them. The most promising candidates for testing include combinations of NSAID (COX2 inhibitors and aspirin), anticoagulant, and lipid-lowering drugs along with antihypertensive drugs such as angiotensin II receptor, calcium channel, or beta blockers, or ACE inhibitors. Combinations of NSAID, anticoagulant, and lipid-lowering drugs that also include more than one antihypertensive drug and, where not contraindicated, estrogen/progestin would be of particular interest.

The current study has two main limitations. First, the drugs in the databases are not identified individually but by category. Many categories include drugs of different classes, which may affect neurodegeneration in different ways. Second, the ideal desired endpoints for the clinical trials would be reductions in biomarkers of neurodegeneration rather than improvements in cognitive ability, but the databases so far exclude biomarkers. AIs trained via ML on large datasets from databases that identify drugs individually and that include biomarkers of neurodegeneration could be used to make better predictions concerning combinations of repurposed drugs that would be effective in treating AD specifically and neurodegeneration more generally.

## ACKNOWLEDGEMENTS

A grant from the Alzheimer Disease Research Fund, administered by the Illinois Department of Public Health (IDPH), supported this research. The IDPH was not involved in any phase of the project. Access to data was provided by the Rush Alzheimer’s Disease Center (RADC) of the Rush University Medical Center in Chicago, IL. The data provided by RADC comes from the participants in the Religious Orders Study and Rush Memory and Aging Project (ROSMAP). ROSMAP is supported by National Institute on Aging grants P30AG10161, R01AG15819, and R01AG17917, and by the Illinois Department of Public Health. Access to data was also provided by the National Alzheimer’s Coordinating Center (NACC). NACC is funded by the National Institute on Aging grant UO1 AG016976, and located in the Department of Epidemiology at the University of Washington School of Public Health.

