## Supplementary Material for "Predicting the potency of anti-Alzheimer drug combinations using machine learning"

*Supplementary Material for:*
title: **Predicting the potency of anti-Alzheimer drug combinations using machine learning**
author: Thomas J Anastasio

This supplement contains items relevant to the above-referenced article. The article describes the results of studies using machine learning to extract the knowledge contained in two Alzheimer Disease (AD) databases, and then using that knowledge to predict combinations of drugs that could be effective in AD treatment. The study was entirely computational. All computations were performed using MATLAB™ running on Intel Core i5 computers, operating in parallel over all four cores. This supplement contains information that supports the narrative developed in the main text. See main text for references.

**Supplement Table of Contents**

| Item | Page |
|---|---|
| Supplementary Note N1 | 2 |
| Supplementary Note N2 | 3 |
| Supplementary Note N3 | 3 |
| Supplementary Table T1 | 5 |
| Supplementary Note N4 | 6 |
| Supplementary Figure S1 | 6 |
| Supplementary Figure S2 | 7 |
| Supplementary Figure S3 | 8 |
| Supplementary Figure S4 | 9 |
| Supplementary Figure S5 | 10 |
| Supplementary Figure S6 | 11 |
| Supplementary Figure S7 | 12 |

**Supplementary Note N1**

The two AD databases providing data for this study were assembled by the Rush Alzheimer Disease Center (RADC) and the National Alzheimer Coordinating Center (NACC). Both of them contain data on elderly participants and focus on AD and other dementias, but they are distinct nevertheless. NACC contains data from the 29 Alzheimer's Disease Centers (ADCs) that are funded by the National Institute on Aging. ADCs are largely tertiary-care dementia centers, and most of the participants already suffered AD or another dementing disease. NACC grew from the Minimal Data Set (MDS) initiated at Rush University in 1997, and this format was used to compile data from various centers in 1998. As its name implies, the MDS was a brief, 50-item description of ADC participants. In 2002 NACC created an improved format for standardized, longitudinal clinical evaluations known as the Uniform Data Set (UDS), and a new version of the UDS was implemented in 2015. As of 2019, the NACC UDS contains data from over 42,000 participants, and is one of the largest dementia databases in the world.

RADC records data from the Religious Orders Study (ROS) and the Memory and Aging Project (MAP), initiated in 1994 and 1997, respectively. The two are collectively known as ROSMAP, and are included together in the RADC database, which now has over 3300 participants. ROSMAP is distinct from NACC in several ways. The most important distinction is that ROSMAP is focused on aging rather than on dementia specifically. Also, ROSMAP is community based rather than clinically based. Whereas most NACC study participants entered as ADC patients and already suffered dementia, ROSMAP participants entered as members of a community whether or not they already suffered dementia. Other differences are that ROSMAP requires an agreement for organ donation but NACC does not, while NACC requires data in the UDS format but ROSMAP does not. ROSMAP data are not recorded in UDS format.

Despite their differences, ROSMAP and NACC share essential features. Both are based on US cohorts and on studies funded by the National Institute on Aging. Both are large, highly regarded, and used by researchers worldwide. Both are concerned with dementia generally and AD specifically. Both record similar data fields: age, demographics (sex, race, etc), tobacco/alcohol use, comorbidities, drug use, and the scores on many tests of cognitive function. Because NACC encompasses data from 29 ADCs it has many redundant fields; these were removed as part of the preprocessing required to make the NACC dataset suitable for machine learning (see main text). Despite that, NACC has fewer drug fields than ROSMAP. Nevertheless, the NACC and ROSMAP databases have 17 drug fields in common. The 100K+ possible combinations of those drugs provide an extensive testbed for the comparison of predictions that result from machine learning on the datasets contained by the ROSMAP and NACC databases.

ROSMAP data is not included in NACC but RADC is one of the 29 ADCs, and it contributed early on to NACC. Specifically, about 1.4% of the NACC dataset is legacy RADC data. All RADC data were removed from the NACC dataset for this study. The two datasets analyzed in this study are entirely independent.

Age is the main AD risk factor, so it was of critical importance in this study for machines to learn how age was related to all the other variables. Machine learning was carried out on age-advancing sequences of at least ten database entries. Though ROSMAP includes many fewer participants than NACC (3300 compared with 42,000; more than an order of magnitude fewer) it is more thorough, so the percentage of participants with ten or more entries was much higher for ROSMAP than for NACC. ROSMAP has 1086 sequences of 10 or more entries for a total of 15,689 entries, while NACC has 2348 sequences of 10 or more entries for a total of 26,434 entries. Restriction to age-advancing sequences of ten or more entries not only enriched the machine-learning dataset with long sequences, but it also served to even out the numbers of entries in the two datasets. The number of machine-learning training iterations was twice the number of sequences for either the ROSMAP or NACC datasets.

**Supplementary Note N2**

The neural networks (NNs) considered as candidates for the optimal NN for this application had three layers: input, internal, and output. The numbers of units in the input and output layers are fixed by the numbers of inputs and desired outputs in the dataset used to train the NN. The internal layer, in contrast, can have many more degrees of freedom. The internal layer of the candidate NNs considered for this study was composed of specialized, compound units known as long short-term memory units (LSTMs). The LSTMs received not only forward connections from input units but also recurrent connections from the other LSTMs in the internal layer. The recurrent connections formed many feedback loops between the LSTMs. These external loops could work in conjunction with internal loops within LSTMs.

LSTMs have a complex structure. Each LSTM contains an internal feedback loop. In addition, each LSTM incorporates five nonlinear units: an input and output unit and three gate units, one each that gates (modulates) the input, output, or internal feedback loop. All five units in each LSTM receive both forward and recurrent connections. Additionally, the three gate units receive peephole connections from the internal feedback loop. Thus, recurrent networks of fully featured LSTMs have eight distinguishing characteristics: external loops between LSTMs, internal loops within LSTMs, three gates, and three peepholes. Altogether there are 256 combination of these 8 characteristics. An NN with any one of these 256 combinations is viable in that it can be trained to improve its performance over the dataset. The best configuration for a given application may be a simplification of a recurrent network of fully featured LSTMs. That was the case for this study: for both the ROSMAP and NACC datasets, the best network had only one of these eight characteristics: the input gate (see main text and Supplementary Note N3).

**Supplementary Note N3**

The genetic algorithm (GA) operates on a population of artificial chromosomes, where each chromosome contains multiple genes and each gene encodes the value of a property to be optimized. The GA involves selection from the population according to fitness, and generation of a new population through variation via random recombination and random mutation of selected chromosomes. The chromosomes in the initial population are random, but the GA improves the fitness of the population as it moves it through the generations. The solution of a GA run is the fittest chromosome in the final generation. Because of the randomness inherent in the GA, the best solution represent the consensus over multiple GA runs.

The GA was used to optimize the characteristics of neural network (NNs) used to extract knowledge from the two AD databases (RADC or NACC). Because the optimal parameters of the machine learning (ML) algorithms used to train NNs can depend on their configurations, the GA optimized ML parameters and NN characteristics simultaneously. The NNs considered as candidates for this study had three layers: input, internal, and output, which had forward connections between them. The internal layer was composed of specialized neural units known as long short-term memory units (LSTMs), which could be interconnected with recurrent connections (see Supplementary Note N2). The ML algorithm used to train recurrent networks with LSTMs is backpropagation through time (BPTT). The version of BPTT used here had three parameters: starting and ending learning rate and fixed momentum (see main text).

To use the GA to optimize NN characteristics and ML parameters simultaneously, artificial chromosomes were configured with twelve genes. The first gene specified the presence or absence of recurrent loops between LSTMs. The second through eighth genes specified the presence or absence of the LSTM features: internal loops, each of the three gates, and each of the three peepholes (see Supplementary Note N2). The ninth gene specified the number of LSTMs in the network. The tenth through twelfth genes specified the ML parameters: starting and ending learning rate and fixed momentum. The NN with characteristics and ML parameters as specified by each chromosome was randomized, retrained, and retested ten times and the generalization errors were averaged (see main text).

The standard GA (with genetic operators of crossover and mutation) operated on a population of 100 chromosomes, each with 12 genes encoding NN and ML properties as outlined above. The GA moved the population through the generations until the average decrease in generalization error over several succeeding generations failed to exceed a tolerance of 1e–3. This always occurred within 100 generations. The GA was implemented in parallel using the MATLAB `ga` command, with the maximal number of generations set at 100. Ten GA runs were performed separately for the ROSMAP and NACC datasets, producing twenty runs altogether. The results of these twenty runs are shown in Supplementary Table T1.

The GA results were consistent between the two datasets and the consensus was obvious. The best generalizing NNs use neither external nor internal memory loops (Ext Mem or Int Mem, columns 1 and 2 of Supplementary Table T1). This is less surprising than it may at first appear. The external and internal loops mediate memory in the network. Because age is included as an input, the network does not need to infer age from its previous unit activations, rendering memory inessential. Because the networks do not make use of the internal memory, the forget gate, which modulates the internal memory, and the three peepholes, which provide views onto the internal memory for the three gates, are not relevant (NR, columns 5 through 8 in Supplementary Table T1).

The best generalizing NNs use either the input gate or the output gate but not both (In Gate or Out Gate, columns 4 and 5 of Supplementary Table T1). A minority use neither. Slightly more networks use the input gate rather than the output gate so the consensus network would use the input gate only, but there is functionally little difference whether the input or the output gate is used to modulate the response before it is transmitted from the LSTM to the output layer. The average number of LSTMs in the best generalizing NNs was about 80, while the average starting and ending learning rates and the fixed momentum were 0.0600, 0.0006, and 0.0002, respectively. All drug combination potency predictions were derived from trained NNs having structure and ML parameters as specified by the consensus chromosome.

| | Ext Mem | Int Mem | In Gate | Out Gate | Mem Gate | In Peep | Out Peep | Mem Peep | Num LSTM | LR Start | LR End | Fixed Mom |
|---|---|---|---|---|---|---|---|---|---|---|---|---|
| ROS MAP | 0 | 0 | 1 | 0 | NR | NR | NR | NR | 90 | 0.0622 | 0.0006 | 0.0002 |
| | 0 | 0 | 1 | 0 | NR | NR | NR | NR | 69 | 0.0580 | 0.0004 | 0.0001 |
| | 0 | 0 | 0 | 1 | NR | NR | NR | NR | 90 | 0.0756 | 0.0007 | 0.0002 |
| | 0 | 0 | 1 | 0 | NR | NR | NR | NR | 29 | 0.0691 | 0.0009 | 0.0003 |
| | 0 | 0 | 1 | 0 | NR | NR | NR | NR | 83 | 0.0616 | 0.0006 | 0.0000 |
| | 0 | 0 | 1 | 0 | NR | NR | NR | NR | 87 | 0.0574 | 0.0010 | 0.0003 |
| | 0 | 0 | 0 | 1 | NR | NR | NR | NR | 90 | 0.0770 | 0.0007 | 0.0002 |
| | 0 | 0 | 1 | 0 | NR | NR | NR | NR | 82 | 0.0838 | 0.0007 | 0.0004 |
| | 0 | 0 | 0 | 1 | NR | NR | NR | NR | 96 | 0.0686 | 0.0008 | 0.0001 |
| | 0 | 0 | 0 | 1 | NR | NR | NR | NR | 81 | 0.0845 | 0.0007 | 0.0001 |
| NACC | 1 | 0 | 0 | 1 | NR | NR | NR | NR | 98 | 0.0557 | 0.0003 | 0.0003 |
| | 0 | 0 | 0 | 0 | NR | NR | NR | NR | 61 | 0.0297 | 0.0004 | 0.0001 |
| | 0 | 0 | 0 | 0 | NR | NR | NR | NR | 97 | 0.0471 | 0.0008 | 0.0003 |
| | 0 | 0 | 0 | 1 | NR | NR | NR | NR | 94 | 0.0938 | 0.0004 | 0.0004 |
| | 0 | 0 | 1 | 0 | NR | NR | NR | NR | 98 | 0.0917 | 0.0006 | 0.0002 |
| | 0 | 0 | 0 | 0 | NR | NR | NR | NR | 95 | 0.0341 | 0.0002 | 0.0001 |
| | 0 | 0 | 1 | 0 | NR | NR | NR | NR | 87 | 0.0704 | 0.0005 | 0.0002 |
| | 0 | 0 | 0 | 0 | NR | NR | NR | NR | 81 | 0.0350 | 0.0009 | 0.0002 |
| | 0 | 0 | 0 | 0 | NR | NR | NR | NR | 81 | 0.0305 | 0.0010 | 0.0001 |
| | 0 | 0 | 0 | 0 | NR | NR | NR | NR | 81 | 0.0302 | 0.0004 | 0.0002 |
| Cons | 0 | 0 | 1 | 0 | 0 | 0 | 0 | 0 | 80 | 0.0600 | 0.0006 | 0.0002 |

**Supplementary Table T1** Results of optimization of NN structure and ML parameters. All optimizations were carried out using the genetic algorithm (GA). The table shows the best chromosome from each of ten GA runs on either the ROSMAP or the NACC dataset. Each chromosome bore twelve genes. The data types were binary for the first eight genes (external and internal memory; input, output, and forget gate; and input, output, and memory peephole), integer for the ninth gene (number of LSTMs), and real for the rest (starting and ending learning rate, and fixed momentum). The memory gate and the three peepholes are not relevant (NR) because the internal memory is absent in all of the best chromosomes. The consensus chromosome (Cons) is based on the averages of the best chromosomes in round numbers.

**Supplementary Note N4**

Creation of the standard input occurred in seven steps. First, an age-advancing sequence of 100 ages from 50 to 110 years was generated (see main text). Second, all input data in either database were ordered according to the age of the participant at the time of each database entry. These ages were not uniformly spaced. Third, a vector of uniformly spaced ages, one for every database entry, was generated. Fourth, all of the input points in each data field were connected via linear interpolation. Fifth, the interpolated data were resampled at the ages in the vector of uniformly spaced ages. Sixth, the resampled, interpolated data were digitally low-pass filtered below the antialiasing frequency (Nyquist frequency) corresponding to the age-advancing sequence of 100 ages. Seventh, the filtered input in each data field was resampled at the 100 ages in the age-advancing sequence. The fourth through seventh steps were accomplished using the MATLAB `resample` command.

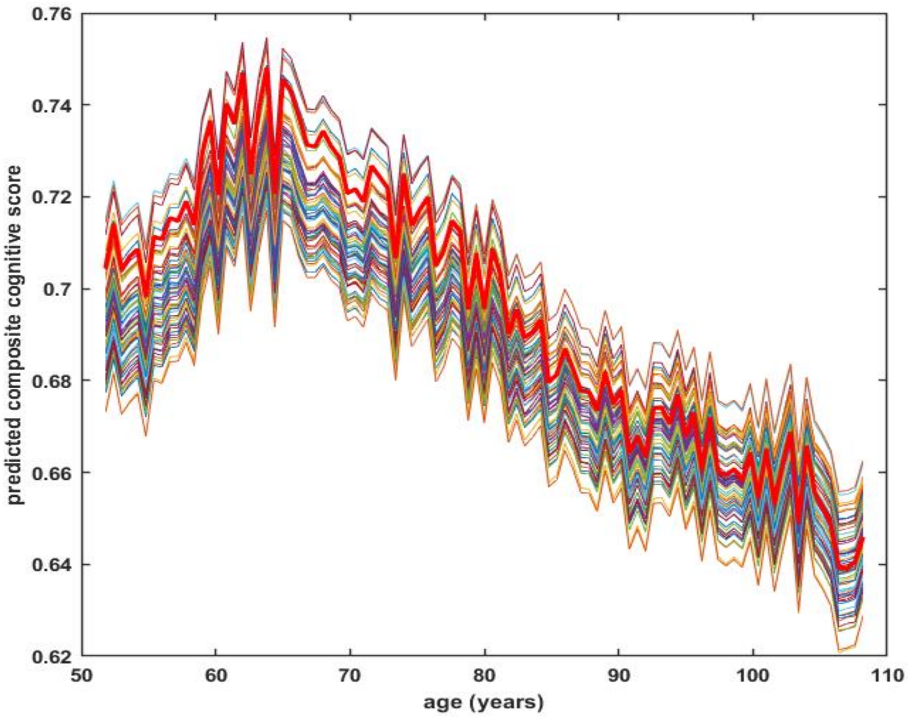

**Supplementary Figure S1** The combined cognitive scores as predicted by a single NN for each age in the age-advancing sequence for 65 representative drug combinations (every 2000$^{th}$ combination selected from the full set of 131,072 combinations of 17 drugs). Each output unit represented the score of a different cognitive test, so the combined cognitive score was the average over the output unit activations. The combined cognitive score in the no-drug case is shown as a heavy red line. This NN was trained on the NACC dataset. The results for NNs trained on the ROSMAP dataset are similar (see Figure 2 of main text). For both ROSMAP- and NACC-trained NNs, many drug combinations are associated with higher cognitive scores than for no-drugs over most or all of the age range. The main difference between the ROSMAP- or NACC-predictions is that the predicted scores rise in the decade from the 50's to the 60's for NACC but not for ROSMAP. This difference likely results because the NACC dataset consists mainly of AD patient entries, and many patients in their 50's likely had early onset AD. In contrast, the ROSMAP database consists of elderly participant entries, and most of the participants were not demented when they entered the Religious Orders Study or the Rush Memory and Aging Project, which provided the data for the ROSMAP database (see Supplementary Note N1).

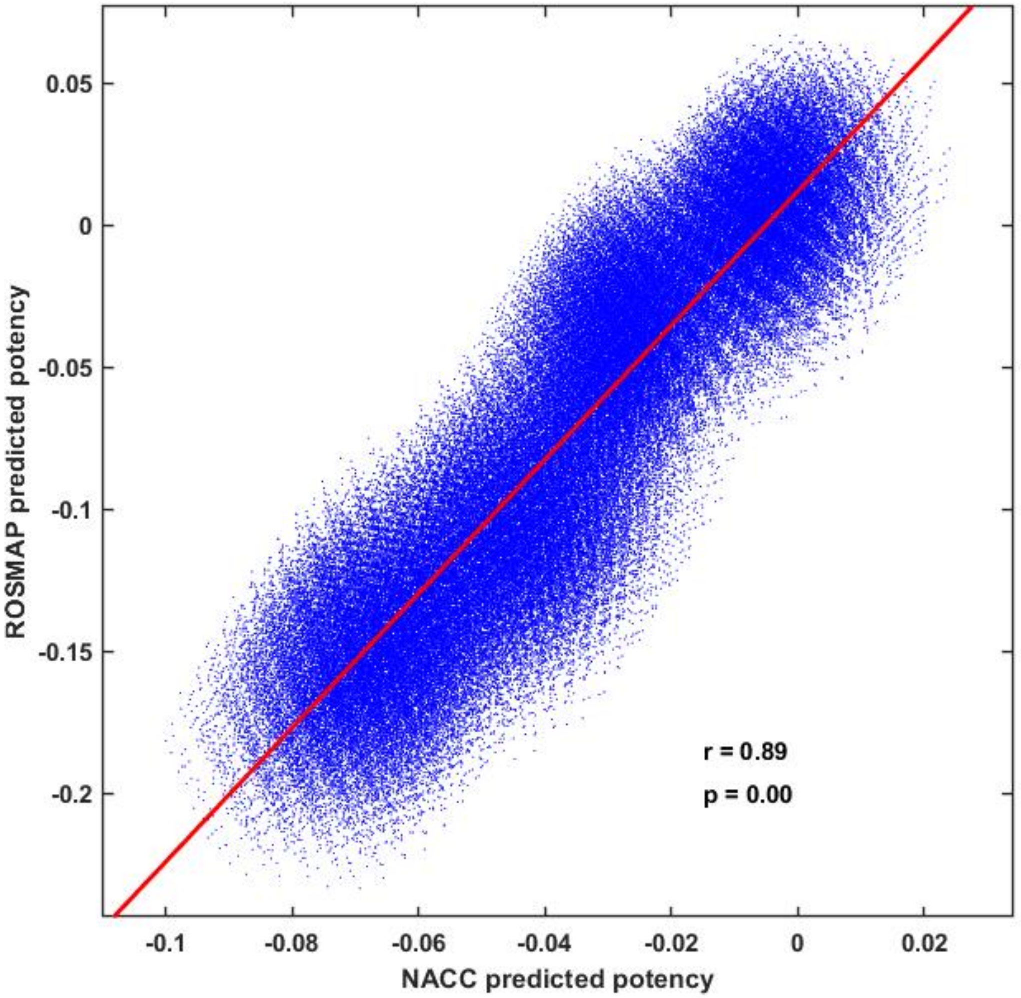

**Supplementary Figure S2** Regressing ROSMAP on NACC predicted potencies, rather than the other way around. Each blue dot locates one of the 131,072 combinations of 17 drugs according to its ROSMAP versus NACC predicted potency. To fit a line to the ROSMAP and NACC predicted potency data, it was necessary to declare one the independent variable (x-axis) and the other the dependent variable (y-axis), but that selection is arbitrary in this case because the ROSMAP and NACC datasets are completely independent of one another. Figure 3A of the main text shows the line resulting from the regression of NACC against ROSMAP. Shown here is the line resulting from the regression of ROSMAP against NACC. Although the slopes and intercepts of the two lines are different, the orderings of drug combinations according to their projections onto the regression line are identical (see also Supplementary Figure S5).

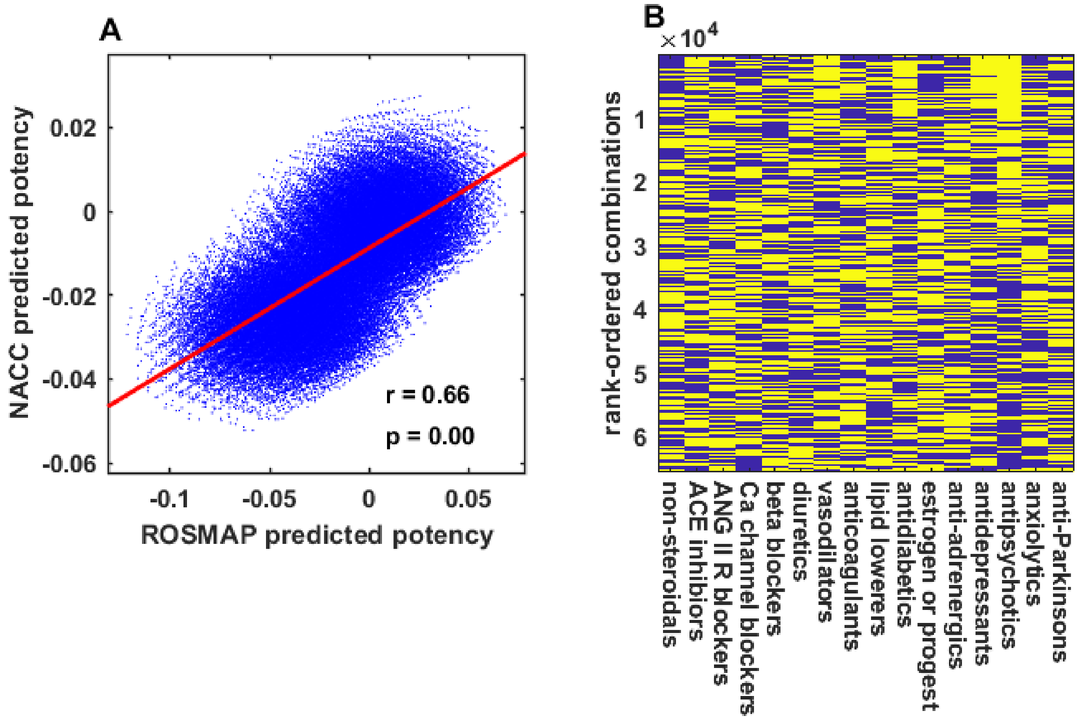

**Supplementary Figure S3** Predicting beneficial drug combinations jointly from NNs trained separately on the ROSMAP or NACC datasets for the 65,536 combinations of 16 drugs that exclude the anti-Alzheimer drugs. (A) Drug combination potencies predicted by NNs trained on the ROSMAP or NACC datasets are strongly correlated. Each blue dot represents one drug combination, located by its ROSMAP and NACC predicted potency (r is the correlation coefficient, p is the probability that the correlation occurred by chance). (B) All 65,536 drug combinations are ranked according to predicted potency (top is highest) and displayed as a heat map (yellow, drug present; blue, drug absent). The most beneficial drug combinations include antipsychotic and antidepressant drugs.

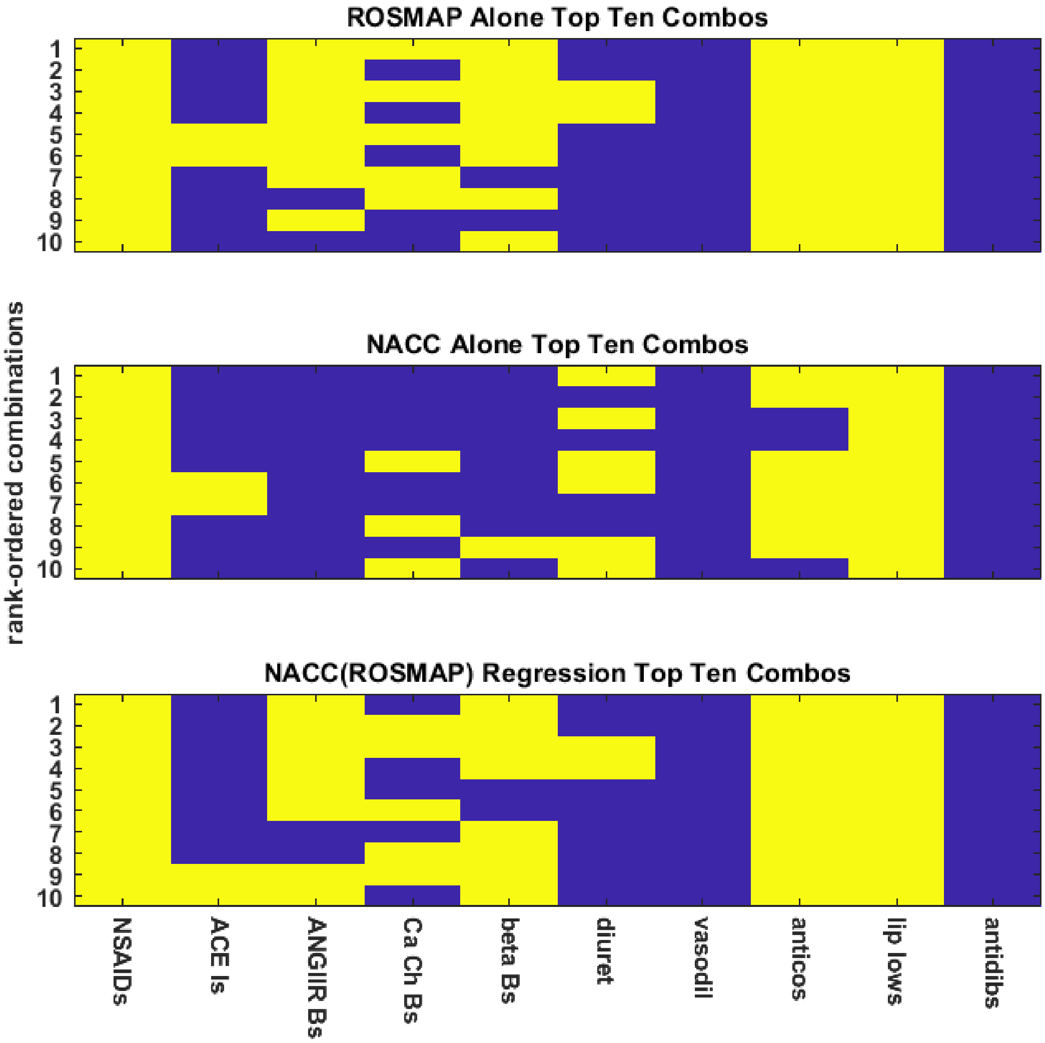

**Supplementary Figure S4** Using either ROSMAP or NACC predictions alone to determine the ten best among the 1024 combinations of the 10 drugs that do not include estrogen/progestin or that target cognitive ability or mood. The ROSMAP alone and the NACC alone ten-best rankings are compared with the ranking based on the regression line giving NACC as a function of ROSMAP (NACC(ROSMAP)); note that the ranking based on ROSMAP(NACC) is identical; see Supplementary Figure S5). The ROSMAP alone and NACC alone top-ten combinations are similar to the NACC(ROSMAP) top-ten in that they all include NSAID and lipid-lowering drugs, and most of them include antihypertensive and anticoagulant drugs. The drug category labels are abbreviations of the labels shown in Supplementary Figure S3.

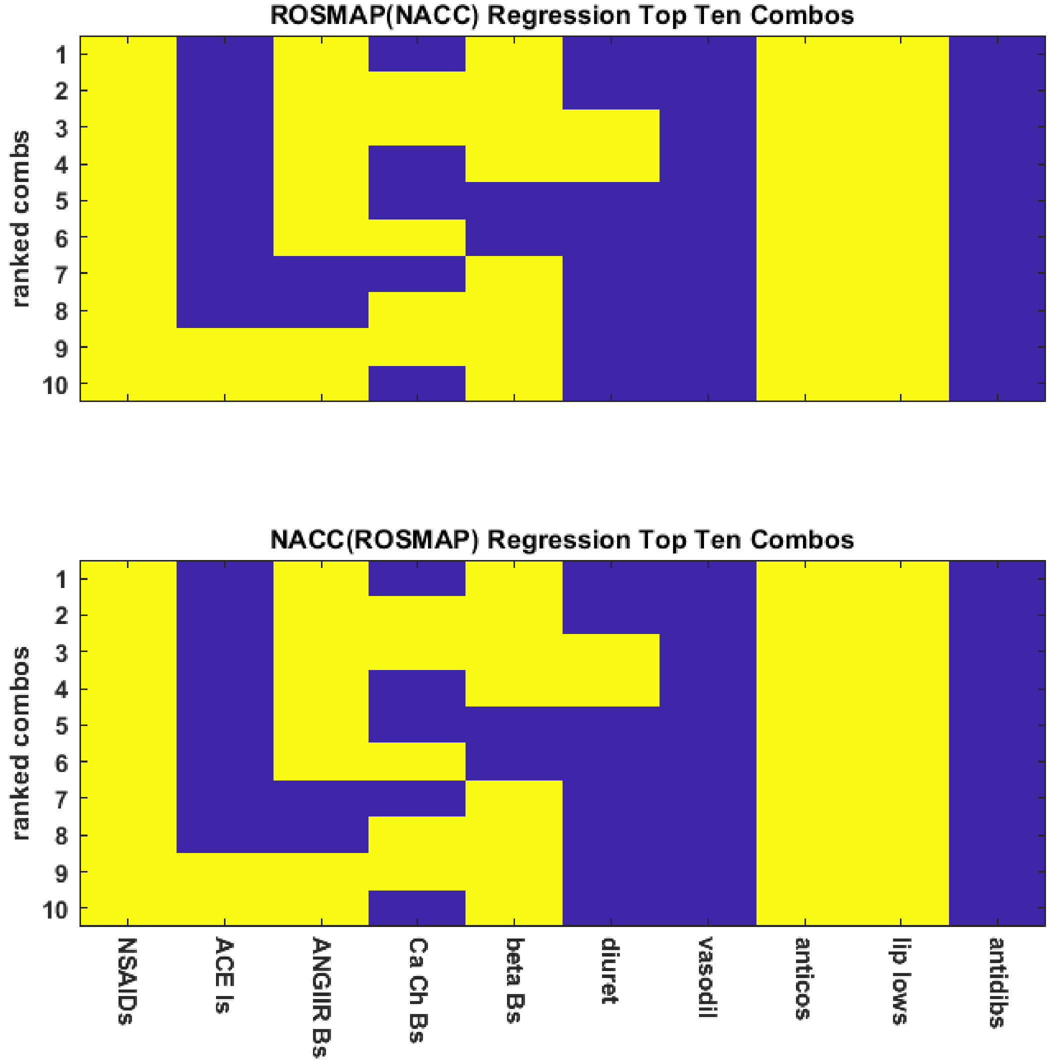

**Supplementary Figure S5** Drug combination predicted potency rankings based on projections onto the regression line giving the NACC prediction as a function of the ROSMAP prediction or vice-versa are identical. The drug combination potencies predicted by NNs trained either on the ROSMAP or NACC datasets are highly significantly correlated (see main text), and joint rankings in terms of linear regression proceed most naturally from this strong correlation. In the main text, the joint determination was made in terms of the projections of the ROSMAP and NACC prediction points onto the line resulting from the regression of the NACC onto the ROSMAP predictions (NACC(ROSMAP)). Because correlation is symmetric, rankings based on NACC(ROSMAP) or ROSMAP(NACC) regressions are identical, as shown here for the top-ten jointly determined combinations. The drug category labels are abbreviations of the labels shown in Supplementary Figure S3.

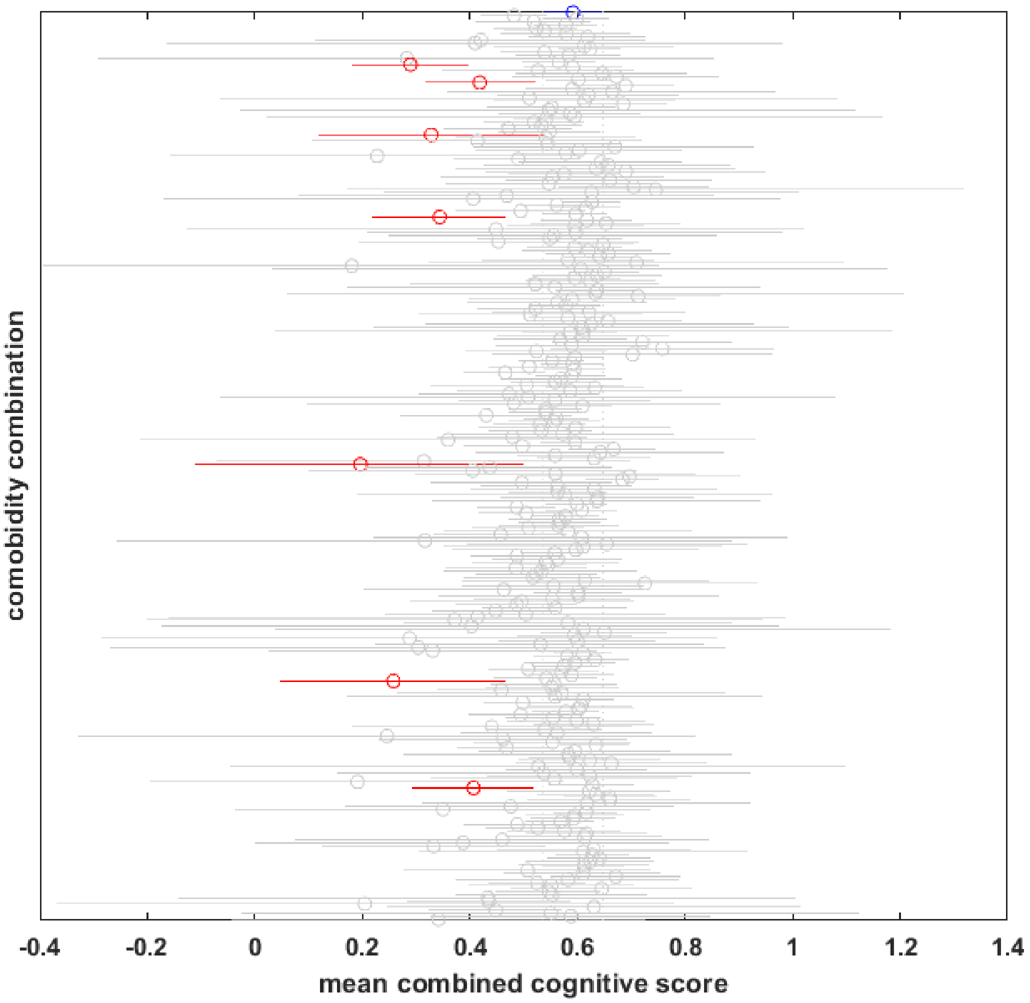

**Supplementary Figure S6** Mean combined cognitive score of ROSMAP participants is not better if they suffer comorbidities. The ROSMAP database has nine comorbidity fields: hypertension, cancer, diabetes, head injury, thyroid disease, congestive heart failure, vascular disease, heart attack, and stroke. Actual ROSMAP participants reported having 298 of the 512 possible combinations of those 9 comorbidities. The blue circle and line at the top of the plot shows the mean and standard error of the cognitive scores of ROSMAP participants with no reported comorbidities. The other circles and lines show the mean and standard error of the cognitive scores of ROSMAP participants with one or more reported comorbidities in the remaining 297 combinations. The red circles show the seven comorbidity combinations associated with mean combined cognitive scores that are significantly lower than the mean associated with no comorbidities. The gray circles show the 290 comorbidity combinations associated with mean combined cognitive scores that are not significantly different from the mean associated with no comorbidities. The statistics were computed using the Bonferroni correction for multiple comparisons.

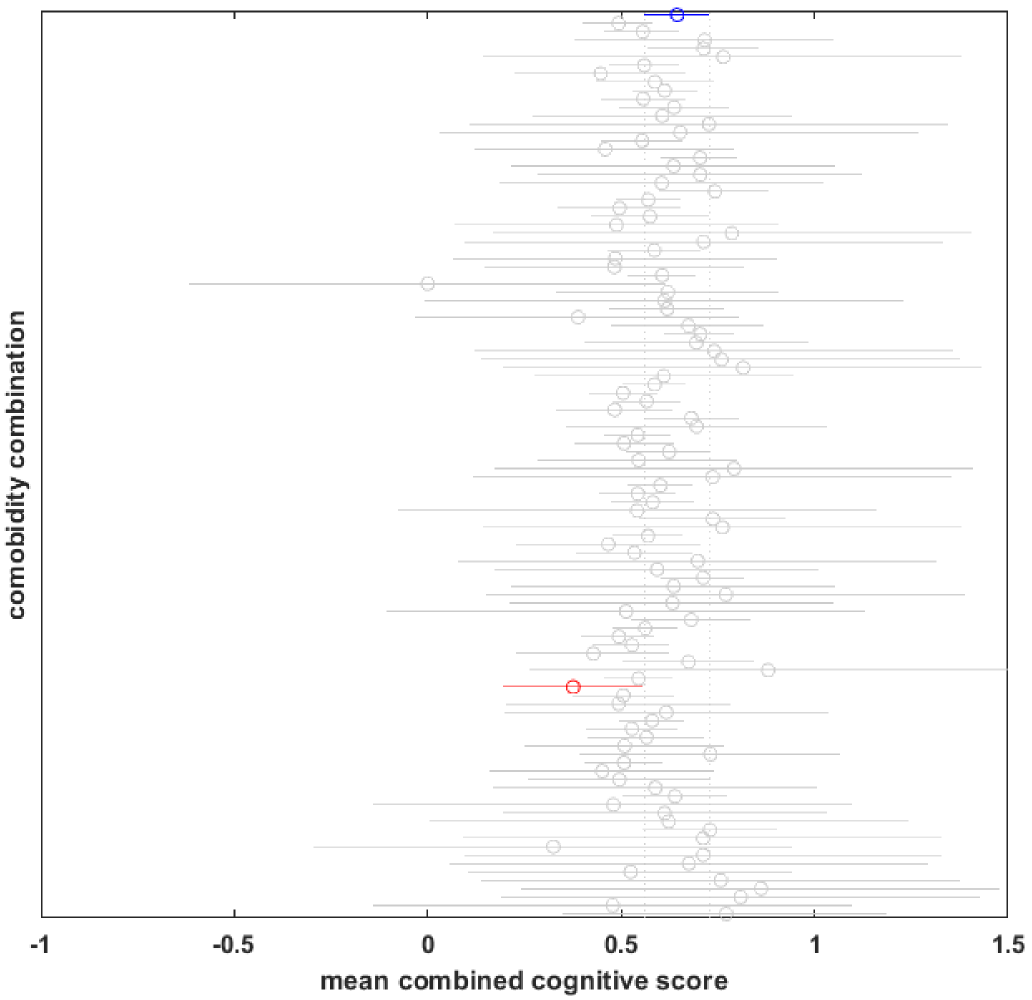

**Supplementary Figure S7** Mean combined cognitive score of NACC participants is not better if they suffer comorbidities. The NACC database has 101 comorbidity fields. Nine of them match or are closely analogous to the nine ROSMAP comorbidity fields. Actual NACC participants reported having 110 of the 512 possible combinations of those 9 comorbidities. The blue circle and line at the top of the plot shows the mean and standard error of the cognitive scores of NACC participants with no reported comorbidities. The other circles and lines show the mean and standard error of the cognitive scores of NACC participants with one or more reported comorbidities in the remaining 109 combinations. The red circle shows the single comorbidity combination associated with a mean combined cognitive score that is significantly lower than the mean associated with no comorbidities. The gray circles show the 108 comorbidity combinations associated with means that are not significantly different from the mean associated with no comorbidities. The statistics were computed using the Bonferroni correction for multiple comparisons.
